# METTL9 regulates N1-histidine methylation of zinc transporters to promote tumor growth

**DOI:** 10.1101/2021.04.20.440582

**Authors:** Mengyue Lv, Dan Cao, Liwen Zhang, Chi Hu, Shukai Li, Panrui Zhang, Lianbang Zhu, Xiao Yi, Chaoliang Li, Alin Yang, Zhentao Yang, Yi Zhu, Kaiguang Zhang, Wen Pan

**Author notes:** corresponding authors: Dan Cao,; Wen Pan. These authors contributed equally (Mengyue Lv, Dan Cao).

## Abstract

Methyltransferase like 9 (*Mettl9*) is a member of the methyltransferase like protein family which is characterized by the presence of binding domains for S-adenosyl methionine, (SAM), a co-substrate for methylation reactions. Despite METTL9 is predicted to be a methyltransferase, its enzymatic activity, substrate specificities and biological functions are still poorly characterized. In this study, we revealed a tumor-promoting role for METTL9. We found that deletion of *Mettl9* in tumor cells suppresses tumor growth and elicits potent anti-tumor immunity. Mechanistically, METTL9 is a N1-histidine methyltransferase which methylates the histidine residues of a x-His-x-His (xHxH) motif on the substrates. This motif is found extensively in zinc transporter families SLC39s and SLC30s, particularly in SLC39A7. Deletion of *Mettl9* impairs cytoplasmic zinc homeostasis, resulting in an altered gene expression program with increased endoplasmic reticulum (ER) stress and reduced cell cycle. Mutation of key METTL9 catalyzed methylhistidine residues of SLC39A7 impairs cytoplasmic zinc homeostasis and affects cell growth as well. Notably, *METTL9* expression is increased in some cancer types and its higher expression is associated with worse clinical outcomes, particularly in liver and pancreatic cancer. In summary, our work identified *METTL9* as a potential new oncogene and its mediated methylation is of regulatory importance. Identifying selective and potent small-molecule inhibitors of METTL9 could thus represent novel therapeutic strategy for anti-proliferative cancer drugs.

## Introduction

Protein methylation frequently occurs as a post-transcriptional modification, which refers to the transfer of CH3 group from a S-adenosyl methionine (SAM, AdoMet) donor to specific amino acid residues in the target protein(1). The target residues include lysine, arginine, histidine, glutamate, glutamine, asparagine and etc.(2). For many years, the main interest has been focused on the N-methylation of protein on lysine and arginine residues, particularly in histones(3-5). Recently, a growing body of works has established the importance of non-histone protein methylation(6, 7), primarily on lysine and arginine residues. It’s worth noting that methylation of proteins at non-lysine/arginine residues is also of profound physiological significance in eukaryotes(8).

Histidine can be methylated at either N1 or N3 position of its imidazole ring, yielding the isomers 1-methylhistidine (His(1-me)) or 3-methylhistidine (His(3-me)). Histidine methylation has been known for many years(9), but only a few proteins carrying such modifications have been studied, including the Actin, Rpl3 etc.(10-14). However, recent proteomic studies suggest that more than 13% of all protein methylation events in human methylome was attributed to the modification of protein histidine residues(15). The results imply that the protein histidine methylation is a widespread modification in mammalians. Nevertheless, the protein histidine methylation has been still rarely studied in contemporary biochemistry. The main reasons may be the complexity of molecular mechanisms required in protein methylation, as well as the lack of sensitive methods to detect methylation at histidine residues *in vivo*(2). More importantly, it is largely unknown whether protein histidine methylation is of profound physiological significance in mammalians, as well as the responsible methyltransferases. So far, only very limited histidine specific methyltransferases were identified in mammalians. Hpm1 was first identified as a Rpl3-specific histidine N-methyltransferase in yeast(14). Histidine methylation of Rpl3 in yeast regulates peptide elongation on ribosomes. SETD3 was recently reported as a dual methyltransferase and modifies both histidine and lysine residues, dependent on the pH on the reaction environment(2, 16). When the pH is seven and above, SETD3 acts as an actin-specific histidine N-methyltransferase. Impaired activity of SETD3 results in the depletion of filamentous actin and a loss of cytoskeleton integrity, as well as labor dystocia in mouse(12). Notably, both Hpm1 and SETD3 modify N3-methylhistidine on the substrate protein, while the methyltransferases that modify N1-methylhistidine remain a major knowledge gap in this field.

The methyltransferase like protein family (METTL) is characterized by the presence of binding domains for SAM, a co-substrate for methylation reactions. Recent findings highlight METTL protein as an important family of putative methyltransferase. For example, METTL3 and METTL14(17, 18), as well as METTL16(19), catalyze the formation of m6A (N6-methyladenosine) in mRNA which is of profound significance in multiple biological processes and diseasees(20). METTL10 and METTL12 have been shown to be protein methyltransferase(21-23), whereas METTL14 may methylate DNA(24). However, for some members of METTL family such as METTL9 and etc., it is still unclear whether they are indeed active enzymes and what are their substrates. Therefore, deep understanding of the biological functions of METTL family proteins and their mediated modifications in mammalians remain a major knowledge challenge and need urgently to be addressed.

In our work, we found *Mettl9* promotes tumor growth. Deficiency of *Mettl9* impairs intracellular zinc homeostasis, markedly diminishing tumor cell growth both *in vitro* and *in vivo*. We further identified METTL9 as a N1-histidine methyltransferase, which catalyzed primarily the latter histidine residues of a xHxH motif on the substrates. METTL9 tightly regulates zinc transporter SLC39A7, which contain a high number of zinc-binding and xHxH-enriched sequences. Mutation of the key methylhistidine residues of SLC39A7 impairs cytoplasmic zinc homeostasis and affects cell growth. Considering *METTL9* expression is increased in some cancers and its higher expression is associated with worse clinical outcomes, particularly in liver and pancreatic cancer, strategies targeting *METTL9* might thus have therapeutic potentials in the future.

## Results

### Deletion of *Mettl9* suppresses tumor growth

As *Mettl9* is broadly expressed in various cancer cell lines and its biological functions are unknown, we set out to study *Mettl9*. We first knocked-out *Mettl9* using CRISPR/Cas9 system in RM-1 and MC38 tumor cell lines, which were derived from murine prostate cancer and murine colon adenocarcinoma, respectively. The targeting strategy and the gene knockout efficiency were evaluated (Fig.S1A-1C) and we obtained at least two knockout clones for each tumor cell line. RM-1 cells lack of *Mettl9* (hereafter termed *Mettl9* KO cells) showed significantly reduced growth and colony formation *in vitro* (Fig.1A-C). Similar results were observed in *Mettl9* KO MC38 cells (Fig.S1D). To further characterize whether *Mettl9* ablation compromised cell growth *in vivo*, we used mouse syngeneic tumor models by subcutaneously inoculating wild-type (WT) and *Mettl9* KO RM-1cells into immunodeficient nude mice. Deletion of *Mettl9* significantly inhibited tumor autonomous growth *in vivo* (Fig.1D-F). Considering both immune and non-immune mechanisms might contribute to the tumor phenotype, we next implanted WT and *Mettl9* KO RM-1 cells into immunocompetent C57BL/6 WT mice. Consistent with what we observed in immunodeficient mice, *Mettl9* ablation similarly resulted in compromised tumor growth in immunocompetent mice (Fig.1G-I). Analysis of intratumoral immune cells showed that both the numbers and percentages of CD45^+^ cells were significantly increased in *Mettl9* KO tumors compared to WT tumors (Fig.S1E, F). Among intratumoral immune cells, the percentages of CD4^+^ and CD8^+^ effector T cells were significantly increased (Fig.S1G, H). Collectively, these results indicated that *Mettl9* deletion not only suppresses tumor cell autonomous growth, but also elicits potent anti-tumor immunity to restrain tumor burden.

**Figure 1.**
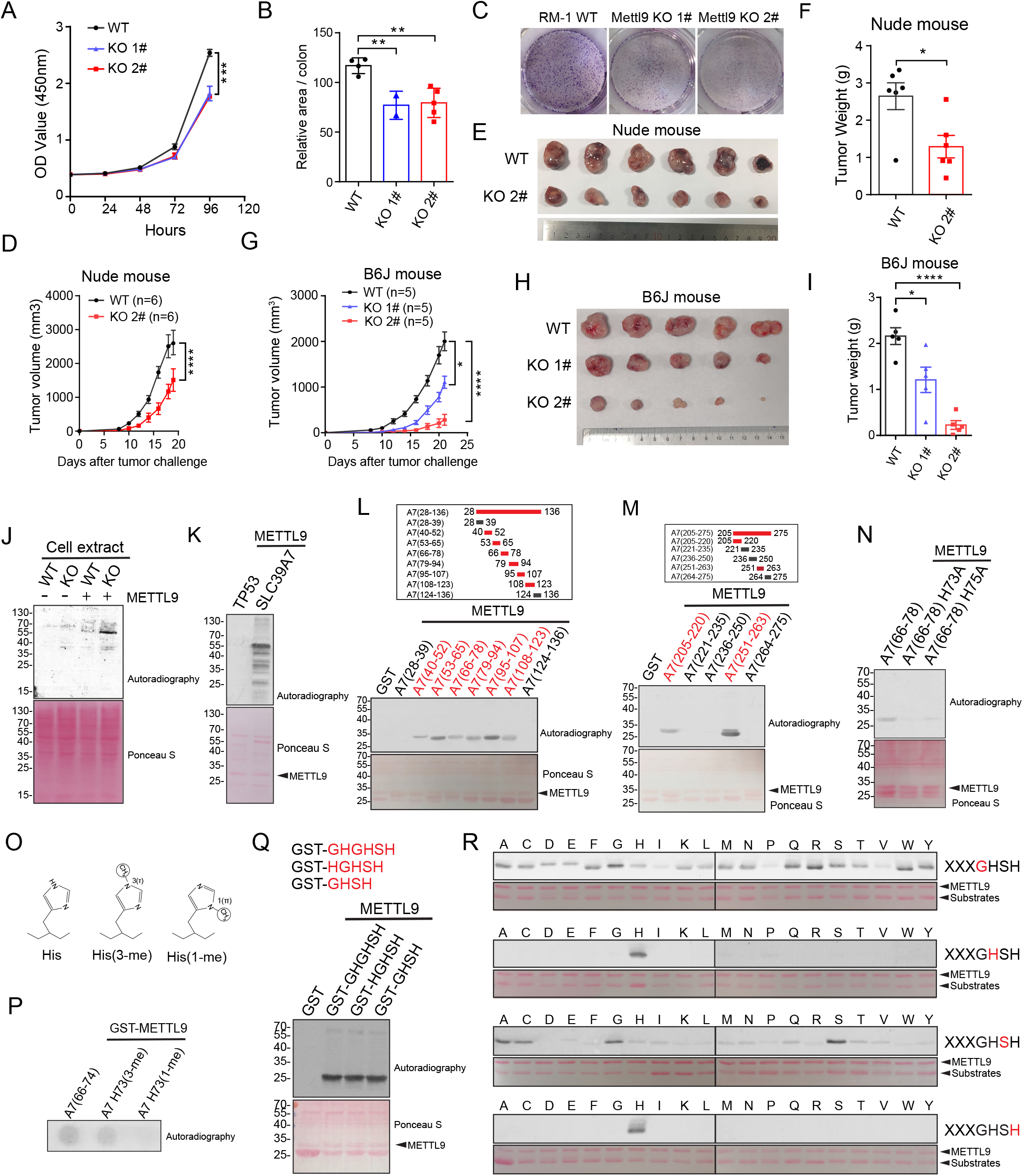
METTL9 is a protein N1-histidine methyltransferase that is required for cell proliferation and tumor growth. **(A, B, C)** Knockout of *Mettl9* in RM-1 tumor cells decreases cell growth. **(A)** Cell growth of wildtype (WT) and two different knockout (KO) clones of RM-1 cells was measured by CCK8 and **(B, C)** colony formation assay. **(D-I)** WT and *Mettl9* KO RM-1 tumor cells were injected into nude mouse and C57B6/J mice. **(D, G)** Tumor growth curves in nude mice (n=6) and C57B6/J mice(n=5). **(E, H)** Pictures of tumors three weeks after tumor cell injection. **(F, I)** Tumor weights for **(E, H). (J-N)** Enzymatically active METTL9 methylates SLC39A7. **(J)** Fluorography showing the activity of recombinant METTL9 on cell extracts from WT and *Mettl9* KO RM-1 cells in the presence of [^3^H]AdoMet. Ponceau S staining for total protein was used as loading control (bottom). **(K)** Fluorography showing recombinant GST-SLC39A7 was methylated by recombinant METTL9 in the presence of [^3^H]AdoMet. GST-P53 was used as a substrate control. **(L, M)** Fine mapping of the METTL9-methylated regions in SLC39A7. The methylated truncate was colored in red. **(N)** Fluorography showing *in vitro* activity of METTL9 on WT and mutated recombinant GST-A7(66-78). **(O)** Histidine (His, left), 3-(τ)-methyl histidine (His(3-me), center); 1-(π)-methyl histidine (His(1-me), right). **(P)** METTL9 generates His73(1-me). *In vitro* methylation reactions with GST-METTL9 on the indicated peptides visualized by autoradiography. **(Q)** Fluorography showing GHSH motif is adequate for METTL9 mediated methylation. *In vitro* methylation assay of METTL9 on recombinant GST-GHGHSH, GST-HGHSH and GST-GHSH. **(R)** *In vitro* activity of METTL9 on recombinant protein arrays. Residue in the GHSH motif (red) was replaced with 20 different amino acids. For all panels, *: *P*< 0.05; **: *P* < 0.01; ***: *P* < 0.001; ****: *P* < 0.0001. Error bars represent S.D. Data are representative of three independent experiments.

### METTL9 is a protein N1-histidine methyltransferase

To explore how METTL9 exerts its function from the molecular and biochemical aspects, we next set out to determine the enzymatic characteristics of METTL9. As we know, METTL9 belongs to the seven-strand (7BS) methyltransferase, characterized by a twisted beta-sheet structure and certain conserved sequence motifs, and these enzymes methylate a wide range of substrates, including small metabolites, lipids, nucleic acids and proteins. We aligned the sequence of METTL9 in different species, and found that METTL9 has conserved sequence motifs that are similar to METTL family members that are responsible for protein methylation (Fig.S2A). We therefore asked first whether METTL9 is a histone methyltransferase. However, purified recombinant mouse METTL9 protein could not methylate a histone substrate (Fig.S3A). Considering the fact that METTL9 protein was localized in the cytoplasm (Fig.S3B), we speculated that METTL9 might be a non-histone methyltransferase. When detecting the activity of METTL9 to methylate proteins in WT and *Mettl9* KO tumor cell lysates in the presence of [^3^H]AdoMet by autoradiography, we observed a substrate protein approximately 55-kDa was methylated (Fig.1J), indicating that METTL9 indeed has enzymatic activity and is a non-histone protein methyltransferase.

To further determine which the abundant substrate protein is likely to be, we detected several interacting proteins of METTL9 as potential substrates. In our previous immunoprecipitation-mass spectrum (IP-MS) experiment, we constructed *Mettl9*-3×flag knock-in RM-1 cell line and identified several interactants of endogenous METTL9 in tumor cells (Fig.S3C). Among them, SLC39A7 is a previously published METTL9 interactant, which is a zinc transporter, nearly 55KD and abundantly expressed in cell lysates (Fig.S3D). We next incubated the recombinant GST-tagged SLC39A7 with recombinant METTL9 in the presence of [^3^H]AdoMet, and found SLC39A7 was indeed an *in vitro* substrate of METTL9 (Fig.1K). To fine-mapping of the methylation sites at SLC39A7, we constructed multiple truncated peptides corresponding to different amino acid sites of the full-length SLC39A7 and narrowed it down to several short peptides, which we confirmed could be methylated by METTL9 *in vitro* (Fig.1L-M, S3E-F). We further synthesized two of these short peptides and incubated them with recombinant METTL9 and AdoMet, followed by acid hydrolysis and analysis of the resulting amino acid by liquid chromatography coupled to MS(LC-MS). The results showed only histidine residues in these peptides were methylated (Fig.S3G, H). We next introduced alanine substitution mutations at the methylated histidine residues, and observed that replacement of histidine abolished METTL9 mediated methylation (Fig.1N). Correspondingly, although METTL9 could not methylate histone itself, it could indeed methylate histone H3 with a 6×His tag (Fig.S4A). All these data confirmed that METTL9 is a protein histidine methyltransferase. As we know, histidine can potentially be methylated on the nitrogen in position 1 (π, N1) or 3 (τ, N3) of the imidazole ring, and endogenous SLC39A7 was identified has physiological methylation on histidine at N1 position(12). We next performed *in vitro* methylation assays on a peptide (residues 66-74 of SLC39A7) that spanned His73 and in which His73 was either unmethylated or methylated at the N1 or N3 position (Fig.1O), and observed that METTL9 methylated the unmodified peptide and the His73(3-me)-containing peptide, but not the His73(1-me)-containing peptide, indicating that METTL9 catalyzes methylation of His73 at the physiologically relevant N1 position (Fig.1P).

To further explore which residues in METTL9 were critical for its enzymatic activity, we mutated several residues in METTL9 potential active sites, motif I and motif II regions (all predicted by I-TASSER) (25-27), which are in a twisted beta-sheet structure of seven-strand(7BS) methyltransferase and are recognized as essential conserved sequence motifs. We incubated WT and mutated recombinant METTL9, as well as 6×His tagged histone H3 protein to perform *in vitro* methylation assay, and defined some key residues that are essential for sustaining METTL9 mediated histidine methylation (Fig.S4B). Collectively, all the above data suggest METTL9 is a protein N1 histidine methyltransferase.

### METTL9 methylates xHxH motifs

To gain insights into the molecular basis of how METTL9 recognizes and methylates a histidine substrate, we analyzed the sequence characteristics of the above fine-mapped methylated short peptides of SLC39A7 and found they all carry a common Gly-His-x-His-x-His (GHxHxH) sequence motif, most of them are GHGHSH. As GHGHSH could be separated as two overlapped GHGH and GHSH, we therefore reasoned that GHxH may be a smaller motif that METTL9 catalyzed (where ‘x’ represents other amino acids). Consistent with this hypothesis, we defined that not necessarily a motif with six amino acid (GHGHSH) is required, but a motif with four amino acid (GHSH) is adequate for METTL9 mediated methylation (Fig.1Q). To further evaluate the contribution of different amino acids in each site of GHxH to the overall histidine methylation level, we used GHSH as a base motif, substituted each residue for 20 different amino acids and generated a total 80 GST-tagged recombinant proteins. By *in vitro* methylation assay, we observed that both the second and the fourth histidine of the motif were indispensable for histidine methylation *in vitro*, and alteration of either totally abolished METTL9 mediated methylation. The amino acid in the third x was preferably Alanine, Cysteine, Glycine, or Serine (ACGS), implying that small and uncharged residues in the middle was necessary for efficient histidine methylation, while the other amino acids introduced at this residue largely impaired the modification. In addition, the first amino acid was relatively tolerant and could be substituted by most of amino acids. However, when it was substituted by Isoleucine, Proline or Valine (IPV), METTL9 mediated histidine modification was significantly reduced (Fig.1R). These data suggest that METTL9 catalyzes the histidine methylation on a motif xHxH, where the first x is preferably not IPV and the second x is preferably ACGS. To further check whether proteins containing physiologically methylated histidine residues within this motif was indeed modified by METTL9, we referred to previously published methylome data and further confirmed that S100A9 and NDUFB3 which carry methylhisitdine at N1 position within the sequence GHSH(1-me) and GHGH(1-me)(28, 29), together with a xHxH containing protein ZnT2 as control, could indeed be methylated by METTL9 (Fig.S5A). This result extended METTL9-mediated methylation on xHxH motif to several physiologically relevant substrates *in vivo*.

Since METTL9 methylates xHxH motif and therefore may have multiple substrates, we next asked what’s the possible biological downstream that METTL9 regulates. To this end, we searched in the mouse proteome using METTL9 preferable recognition motif sequence that we identified above (GHSH, GHAH, GHCH, GHGH), the results showed that the top hits were mainly enriched in zinc transporter families SLC39s and SLC30s (Fig.S5B). Like SLC39A7, most of zinc transporter members contain more than one xHxH motif in their sequences (Fig.S5C), implying that METTL9 might be critical for regulating zinc transporters and affecting cellular zinc concentration. Taken together, our data above indicated that METTL9 is a broad specificity enzyme with a preference for xHxH motifs, and might tightly regulate zinc homeostasis.

### Deletion of *Mettl9* impairs cytoplasmic zinc homeostasis

Zinc transporters tightly regulate cellular zinc homeostasis(30). Controlling appropriate intracellular zinc concentration plays an important role in alleviating cell intrinsic stress and maintain cell growth(31, 32). To investigate whether METTL9 regulates cellular zinc concentrations, we utilized a fluorescent zinc probe (Fluozin-3) to evaluate the free zinc concentration in WT and *Mettl9* KO RM-1 tumor cell lines. In *Mettl9* KO cells, we observed that zinc level was significantly increased and aggregated in cytoplasm, although it is a bit difficult to determine whether this aggregation might be due to the sequestration of zinc in the endoplasmic reticulum or other organelles (Fig.2A). The data support that *Mettl9* indeed plays an important role in regulating of zinc homeostasis in cells.

**Figure 2.**
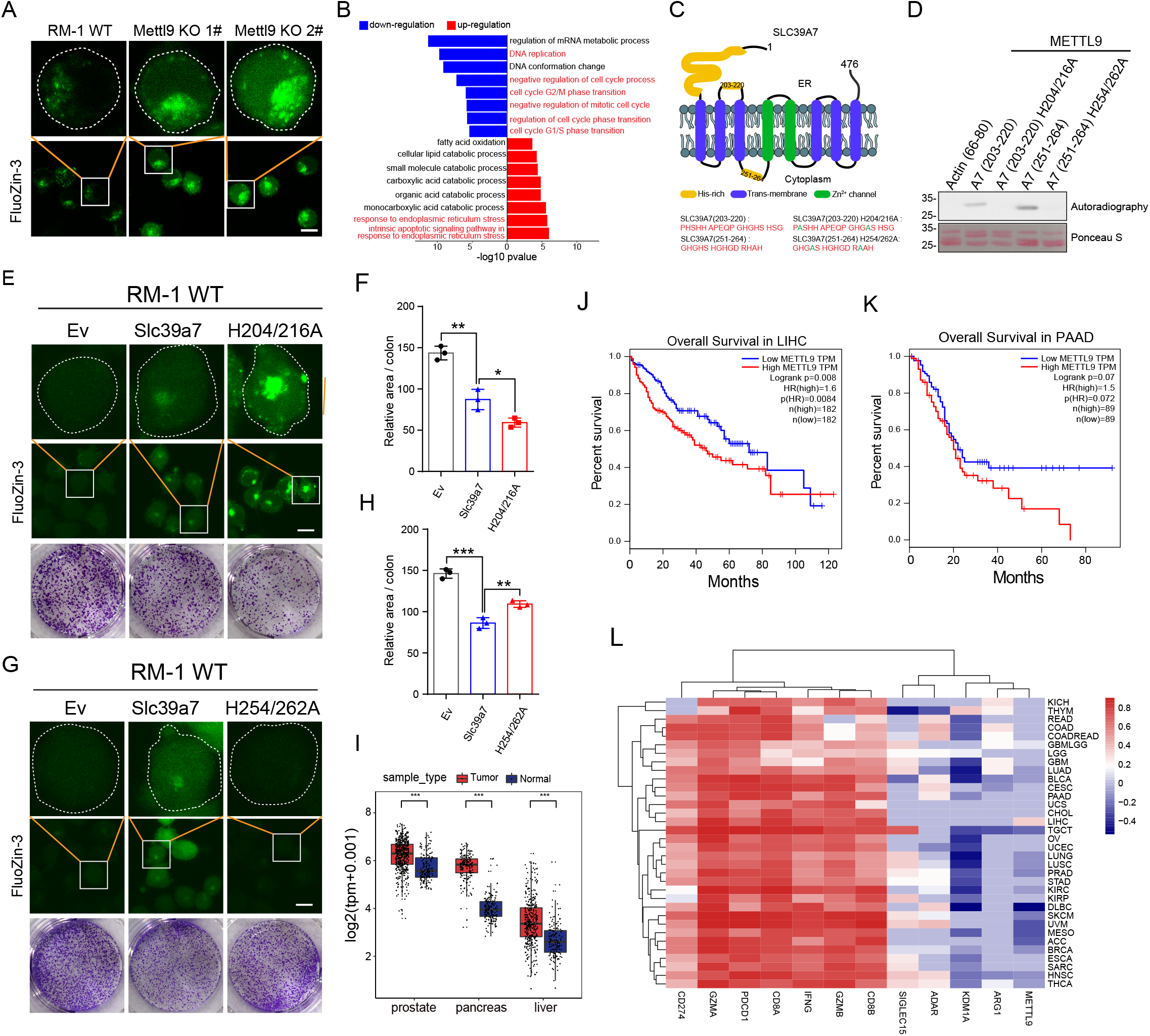
METTL9 catalyzes methylhistidine residues of zinc transporter SLC39A7 and regulates cytoplasmic zinc homeostasis and cell proliferation. **(A)** Fluorescence images of FluoZin-3 (2 μM, 1 h, 25 °C) stained WT RM-1 cell line (left), *Mettl9* KO 1# cell line (middle), *Mettl9* KO 2# cell line (right). Scale bar, 10 μm. **(B)** GO analysis of top pathways from differentially expressed genes in WT and *Mettl9* KO RM-1 tumor cells. **(C)** The predicted structure model of SLC39A7. The number represents the peptide region in SLC39A7. Green color represents mutant residues. **(D)** *In vitro* activity of METTL9 on recombinant GST tagged WT and mutant peptides at the indicated region of SLC39A7. H204/216A, histidine to alanine mutation at 204 and 216 sites; H254/262A, histidine to alanine mutation at 254 and 262 sites. **(E-H)** Overexpression of WT and mutant SLC39A7 in RM-1 WT cells. **(E, G)** Fluorescence images of FluoZin-3 (2 μM, 1 h, 25 °C) stained the indicated groups of BFP-empty vector (EV), BFP-SLC39A7, BFP-SLC39A7 H204/216A and H254/262A mutants. Scale bar,10 μm. **(F, H)** Quantification of colony formation assays in **(E, G)** (see Methods). **(I)** *METTL9* transcript levels were analyzed in cancer and normal tissues from the TCGA database combined with GTEx normal data. PRAD (Prostate adenocarcinoma) PAAD (Pancreatic adenocarcinoma) and LIHC (Liver hepatocellular carcinoma). **(J, K)** The overall survival rates of the *METTL9* high-expression group and the *METTL9* low-expression group in PAAD and LIHC cancer types from the TCGA data. **(L)** Heatmap correlation between several genes expression levels and immune scores calculated by ESTIMATE in tumor tissues among different TCGA cancer datasets. For all panels, *: *P*< 0.05; **: *P* < 0.01; ***: *P* < 0.001; unpaired two-tailed Student’s t-test. Error bars represent S.D. Data are representative of three independent experiments.

As we know, abnormal zinc transport from ER to cytoplasm induces ER stress, and impaired cell proliferation(33, 34). we next performed RNA-seq analysis on WT and *Mettl9* KO RM-1 cells to evaluate the molecular downstream of *Mettl9*. We observed that the transcripts of most zinc transporters did not show obvious expression alteration (data not show). As expected, Gene ontology (GO) enrichment analysis of the differential genes revealed that genes with reduced expression in *Mettl9* KO cells were significantly enriched in GO terms related to ‘Cell cycle’ ‘DNA replication’, whereas those with increased expression in *Mettl9* KO cells were significantly enriched in ‘Endoplasmic Reticulum Stress’ (Fig.2B). Interestingly, it has been reported that *Slc39a7* knockdown (KD) results in abnormal zinc transport from ER to cytoplasm, induced ER stress, and reduced cell proliferation(cite). We reanalyzed the published RNA-seq data from WT and *Slc39a7* KD cells, and GSEA analysis found that gene sets, such as ‘Cell cycle’ or ‘DNA replication’, ‘Homologous recombination’ and ‘Ribosome biogenesis in eukaryotes’, were commonly downregulated both in *Mettl9* KO and *Slc39a7* KD cells compared to their corresponding controls (Fig.S6A-H). Therefore, we reasoned that although METTL9 possibly modify multiple zinc transporters, SLC39A7 appears to be an essential one of them that mediating METTL9 downstream function in cells. Taken together, these data suggest that deletion of *Mettl9* impairs cytoplasmic zinc homeostasis, resulting in an altered gene expression program with increased endoplasmic reticulum (ER) stress and reduced cell cycle.

### METTL9 catalyzed methylhistidine residues of SLC39A7 regulates cytoplasmic zinc homeostasis

We next sought for the key METTL9 catalyzed residues on SLC39A7. According to the previous studies, SLC39A7 harbors highly-conserved histidine-rich regions that potentially contribute to zinc binding and transport (30, 35-37). In these histidine-rich regions, we selected two METTL9 motif-containing fragments (residues 203-220 and residues 251-264 of SLC39A7) for further investigation (Fig.2C), although we could not exclude the possible roles from other histidine sites. We first confirmed that these two SLC39A7 peptides could be methylated by METTL9, whereas peptides with His to Ala mutation impaired METTL9 mediated methylation (Fig.2D). To investigate how these key methylhistidine residues control zinc homeostasis, we constructed two mutated constructs of full-length *Slc39a7*, each one replacing two targeted His residues to Ala to mimic the unmethylated status. We next overexpressed these constructs into RM-1 WT cells and found that H204/216A mutation significantly upregulated zinc aggregation in cytoplasm compared to WT construct controls (Fig.2E), while H254/262A mutation slightly downregulated zinc level (Fig.2G). Consistent with the intracellular zinc levels, we observed that H204/216A mutation significantly downregulated colony formation (Fig.2E, F), while H254/262A mutant slightly upregulated colony formation compared to SLC39A7 WT controls (Fig.2G, H). Although it needs more detailed structure information to elucidate how exactly SLC39A7 mutation affects zinc binding or transport, these data support that some key METTL9 catalyzed methylhistidine residues indeed affect the normal function of zinc transporter SLC39A7, which further regulates cellular zinc homeostasis and cell proliferation.

### *METTL9* shows prognostic potentials in some cancer types

As we know, *METTL9* is broadly expressed in various human cancer cell lines according to Cancer Cell Line Encyclopedia (CCLE) database (Fig. S7A). We next want to evaluate the clinical relevance of *METTL9* in human malignancy. By a comprehensive analysis of the expression data from The Cancer Genome Atlas (TCGA) database and GTEx databases, we found that *METTL9* mRNA expression was significantly increased in some cancers compared with normal tissues, such as pancreatic cancer, liver cancer, and prostate cancer (Fig. 2I). To evaluate the prognostic value of *METTL9* in these cancers, we performed a pan-cancer survival analysis concerning overall survival, using the Cox regression model and log-rank test, for hypothesis test including the Cox proportional hazard ratio. We observed that higher expression of *METTL9* is associated with worse clinical outcomes, particularly in liver cancer and pancreatic cancer (Fig. 2J, K), supporting that *METTL9* may emerge as a prognostic biomarker in these cancer types. Moreover, we applied ESTIMATE computational method to calculate the immune score of different cancer samples from TCGA database. We noticed that *METTL9* gene expression level negatively correlated with immune scores in most cancers, which is consistent with our previous observation that anti-tumor immunity is involved in *Mettl9* KO tumor phenotype (Fig. 2L).

## Discussion

Despite being identified for 70 years; histidine methylation has never been seriously studied(9). Recent proteomic studies suggest that the protein histidine methylation is a widespread modification in mammalians(15). However, it is largely unknown about the functional and (patho)physiological significance of such modification in cells and organisms, as well as the responsible methyltransferases(2). When the first two histidine-specific methyltransferases Hpm1 and SETD3 have been identified, histidine methylation began to attract considerable interest among the scientific community. Here, by using a combination of *in vitro* and *in vivo* approaches, we identified METTL9 as a histidine methyltransferase that specifically catalyzes N1-methylhisitidine formation in distinct protein substrates. We demonstrated that METTL9 promotes tumor growth in cellular and animal models.

During the submission process of our manuscript, Shinkai and Falnes groups first reported METTL9 is a N1 histidine methyltransferase which methylates His-x-His motif in numerous substrates(38). They found METTL9-mediated His(1-me) formation is abundant in mammalian cells and tissues. Our work independently confirmed that METTL9 is a protein N1 histidine methyltransferase and established that METTL9 recognized a xHxH motif in the substrate protein. We demonstrated the contributions of different amino acid in the middle and flanking residues of the recognition motif to histidine methylation, supporting that the x in the middle is preferably Alanine, Cysteine, Glycine, or Serine (ACGS), which is not exactly consistent with the reported ANGST preference in the middle. The major difference may be that we used purified recombinant peptides instead of synthetic peptides as substrates in the methylation reaction. Besides, the modification pattern is more complex when the recognition sequence contains a consecutive xHxH motif. Therefore, further structure analysis on the interface of METTL9 and its modified substrates is urgently needed.

Importantly, our work elucidates the functional and (patho) physiological significance of *Mettl9* and its mediated modification in cells and organisms. Deletion of *Mettl9* inhibits cancer cell proliferation and impaired colony formation capacity, both *in vitro* and *in vivo*. We further revealed that METTL9 tightly regulates zinc transporter families and controls the cellular zinc homeostasis. On the transcriptome level, deletion of *Mettl9* results in an altered gene expression program with increased endoplasmic reticulum (ER) stress and reduced cell cycle. METTL9 mediates methylation of the histidine residues at the zinc transporters, and some key methylhistidine residues are essential for zinc binding and controlling intracellular zinc homeostasis. In line with our findings that loss of *Mettl9* inhibits cancer cell proliferation and impaired colony formation capacity, low expression of METTL9 correlates with increased survival of patients with liver cancer and etc. Besides, *METTL9* gene expression level negatively correlated with immune scores in most cancers. In the future, it will be important to explore the roles of *METTL9* beyond prostate and colon cancer model we used, as *METTL9* has also been found amplified in various tumors.

METTL9 mediated methylation of the histidine residues precisely modulated the affinity of SLC39A7 to bind to zinc. Although multiple histidine residues were shown to be methylated *in vitro*, it is very important to develop sensitive and specific methods for probing protein methylation *in vivo*, and to identify the physiological methylation residues of the substrate. Besides, in regard to substrate proteins carrying more than one methylation residues, it is necessary to combine the structure information of the substrate to better explore the functions of individual methylhistidine site.

Although *METTL9* is abnormally unregulated in various tumor sample, it is unknown how exactly *METTL9* was regulated in context of tumor microenvironment. It would therefore be interesting to explore the upstream regulators of *METTL9*. Especially, whether *METTL9* was modulated by the zinc level in the cytoplasm in a positive-feedback loop, considering METTL9 methylates multiple zinc transporter family members and controls cellular zinc homeostasis. Besides, in a manner analogous to lysine or arginine methylation on histones, we reasoned that METTL9-mediated methylation may be dynamic. It would therefore be important to explore the dynamics of methylation and seek for the potential demethylase that is responsible for erasing METTL9 methylated residues.

Our work identified *METTL9* as a potential new oncogene and its mediated methylation is of regulatory importance. In the future, we believe there will soon be more works published regarding an interesting insight into the physiological roles of METTL9 and other novel protein histidine methyltransferases. Given the recent success in identifying selective and potent small-molecule inhibitors of methyltransferases, histidine methyltransferases such as METTL9 could thus represent novel therapeutic targets for anti-proliferative cancer drugs.

## Materials and methods

### Cell culture

The RM-1, MC38, HEK239T cell line was purchased from American Type Culture Collection (ATCC, Manassas, VA, USA). All cell lines were cultured in high-glucose Dulbecco’s modified Eagle’s medium (H-DMEM: Gibco) supplemented with 10% fetal bovine serum (FBS; Gibco), 100 U/mL penicillin, and 100 ug/ml streptomycin. All cell lines were cultured at 37 °C with 5% CO2.

### Plasmid construction and Peptides

*Slc39a7* (ZIP7) and its fragments and mutants were cloned and inserted into the EcoR I and Xho I sites of pGEX-6P-1. *Slc39a7* and its mutants were inserted into the Xho I and BamH I sites of Plvx-IRES-BFP. *Slc39a7* were inserted into the Nhe I and Xho I sites of pcDNA 3.2 -3’HA. *Mettl9* and its mutants were cloned and inserted into EcoR I and Xho I sites of pGEX-6P-1. *Mettl9* were inserted into EcoR I and BamH I sites of pEGFP-N3. All the peptides GHGHSH, HGHSH, GHSH and GHSH mutants were designed and inserted into EcoR I and Xho I sites of pGEX-6P-1 by annealing. *S100a9* was cloned and inserted into BamH I and Xho I sites of pGEX-6P-1. *ZnT2* and *Ndub3* were cloned and inserted into EcoR I and Xho I sites of pGEX-6P-1. 6×His Histone H3 was a gift from Yong Ding lab in USTC.

The following peptides were synthesized:SLC39A7(66-78): DFHHGHGHTHESI; SLC39A7(95-107): LHHGHSHGHSHDS; SLC39A7(66-74): DFHHGHGHT; SLC39A7(66-74) H73(3-me): DFHHGHG(His(3-me))T; SLC39A7(66-74) H73(1-me): DFHHGHG(His(1-me))T.

### Generation of mouse *Mettl9* KO/KI cell lines

The *Mettl9* gene was disrupted by the CRISPR-Cas9 method using four guide RNA targeted to introns of upstream (sgRNA1: 5′-AAAGATGATGATCGGCCTCA-3′; sgRNA2: 5′-GATGATGATCGGCCTCAGGG-3′) of exon 2 and downstream (sgRNA3: 5′-AGCCATGTGTAGTATCACCG-3′; sgRNA4: 5′-TACAGCATTCTGTACGCCCC-3′) of exon 4 in *Mettl9* were cloned into Bbs I site of HP180 vector (A gift from Hui Yang lab). The guide RNA plasmids then transfected into RM-1 and MC38 cells. Two days after the transfection, GFP positive cells were sorted and single cells were seeded into 96 well plate. The clones were screened by genomic PCR. For the generation of RM-1 knock-in (KI) cells complemented with C-terminally 3×flag-tagged *Mettl9*, plasmids for *Mettl9*-flag donor (pUC19-*Mettl9*-3×flag) and guide RNA plasmids (sgRNA5: 5′-CTCAGACCAGTATAAACACG-3′; sgRNA6: 5′-TCTTCTGGAGGGTCGGTTGC-3′) were co-transfected into RM-1 cells using lipofectamine 2000 transfection reagent. Two days after the transfection, GFP positive cells were sorted, and individual clones were obtained by limiting dilution method. The flag KI clones were screened by genomic PCR and confirmed by Western Blot using anti-flag antibody.

### Western Blots and antibodies

Protein was extracted from the cells with IP buffer (25 mM Tris-HCl pH 7.4, 150 mM NaCl, 1 mM EDTA, 1% NP-40, 5% glycerol) and resolved on SDS–PAGE gels, then transferred to PVDF membranes. The primary antibodies against METTL9 (1:2000, ProteinTech) and β-actin (1:5000, ProteinTech) were used overnight at 4 °C, then washed the PVDF membranes three times with TBST (0.1% Tween-20 in TBS), the membranes was incubated with secondary antibody (1:5000, ProteinTech) for 40 minutes and washed three times with TBST. The membrane was explored by using enhanced chemiluminescence kit (Biosharp) according to the manufacturer’s instructions.

### Real-time RT-PCR analysis

TRNzol (TIANGEN) was used to extract RNA from cells growing on tissue culture dishes. 500 ng RNA were used for following RT-PCR. The HiScript RT SuperMix kit (Vazyme) were used for generating cDNA according to the manufacturer’s instructions. The qPCR was conducted in 384-well PCR microplates (Bio-Rad) in a CFX384 Real-Time System (Bio-Rad). Each well contained a total volume of 10 μl, including 4.2 μl cDNA (concentration ∼2.5 ng/μl), 0.4 μl primers (10 μM), 5.0 μl Fast SYBR Green Master Mix (SYBR premix EX Taq, TaKaRa). The fluorescence was measured at the end of each cycle. Measured data was analyzed using the comparative CT method (ΔΔCT method). Mouse *Rpl13a* prime F: GGGCAGGTTCTGGTATTGGAT, R: GGCTCGGAAATGGTAGGGG; mouse *Mettl9* prime F: CTGGCAGCTCCAGAAGAAGA, R: CCACTTGCCACCTACGTTTT.

### Cell proliferation and colony formation assay

Cell proliferation was quantified using the cell counting Kit-8 (CCK8; APExBIO) according to the manufacturer’s instruction. Cells were seeded in 96-well plates in triplicate at an initial density of 2000 cells per well. After being cultured for 24, 48, 72, and 96 hours, CCK-8 solution in 10 ul was added to each well, and cells were further incubated at 37 °C for 4 hours. Optical density (OD) value was measured spectrophotometrically at 450 nm wavelength.

Cells were plated in 6-well plates (10000 cells per well), and then cultured for 4 days. The overexpression cells were generated by infecting with BFP-Slc39a7 or mutants’ virus. After 24 h, the BFP-positive cells were sorted by flow cytometry. For an assay of colony formation, colonies were fixed with 4% paraformaldehyde for 20 minutes, stained with 0.1% crystal violet for 15 minutes, and then washed with phosphate-buffered saline (PBS) before measurement. The total colony area was measured by image J software. The relative area was calculated by total area divided by original cell number.

### Tumor formation assay

The C57B6/J mice and BALB/c nude mice were feeding under specific pathogen-free conditions in the experimental animal department of USTC. For the *in vivo* tumor formation assays, 1.5×10^5^ RM-1 *Mettl9* KO cells (1#, 2#) or RM-1 wild type cells were subcutaneously injected into the C57B6/J mice (n = 5 for each group) and the BALB/c nude mouse (n = 6 for each group) at 8 weeks of age. After transplantation 8 days, the growth of the tumors was assessed every two days. The mice were sacrificed after a period of about 3 weeks, and the weights of subcutaneous tumors were measured.

### Analysis of immune cell infiltrate in tumor microenvironment

Preparation of single-cell suspension of mouse tumor tissues by mechanical grinding and collagenase digestion. For flow cytometry analysis, all single-cell suspensions were incubated with anti-mouse CD16/CD32 blocking antibody for 15□minutes after thorough filtration and precipitation, stained with fluorescein-conjugated antibodies, multiple washed with PBS. Cells were stained with the following mAbs: anti-CD45, anti-CD4, anti-CD8a, all from Biolegend. Samples were run on a BD flow cytometer and analyzed with the Flowjo Analysis Software (Beckman Coulter, Pasadena, CA). Cell populations were defined as follows: CD45^+^ CD4^+^CD8^-^(CD4^+^ T cells); CD45^+^ CD4^-^CD8^+^ (CD8^+^ T cells).

### IP assay and Mass spectrometry

The 15 cm dish *Mettl9*-3×flag RM-1 cells extracts prepared with IP buffer (25 mM Tris-HCl pH 7.4, 150 mM NaCl, 1 mM EDTA,1% NP-40, 5% glycerol) remain on the ice for 30 minutes. Lysates were cleared by centrifugation at 12000 rpm for 10 minutes and incubated with anti-flag (anti-flag M2 Affinity Gel, Sigma) antibodies covalently attached to agarose 4 hours at 4 °C on a rocking platform. Beads were then collected by centrifugation at 3000 rpm for 5 minutes at 4 °C, extensively washed in lysis buffer, and then the mass spectrum was performed. To select the METTL9-interacting proteins, the PEMs and Sum PEP Score was utilized to do a dot plot and top proteins closest to the upper right corner (PSMs > 2; log2(Sum PEP Score) > 2) was selected. For synthetic peptides methylation detection, after methylated by METTL9, the sample were sent to MS detection.

The processed data were used to search with ProteinDiscovery (version 2.2, Thermo fisher Scientific) against the UniProt mouse filtered organism database or in-house database including the amino acid sequences of synthetic peptides, using the following parameters: enzyme = trypsin; maximum missed cleavages = 1; variable modifications = Acetyl (Protein N-term), Oxidation (M), Methyl (H), Methyl (K), Methyl (R), Propionamide (C); product mass tolerance = ±10 ppm; product mass tolerance = ±0.02 Da (LIFT mode); instrument type = Orbitrap.

### Expression and purification of recombinant proteins

The appropriate pGEX-6P-1 constructs were transformed into the chemically competent BL21-Codon Plus (DE3)-RIPL (Agilent Technologies) E. coli strain. The bacteria were cultured overnight in 3 ml LB medium with 0.1% Ampicillin sodium (BBI) at 37 °C, 250 rpm, and then inoculated into 250 ml LB medium with 0.1% Ampicillin sodium at 37 °C, 250 rpm, after about 5 hours until the OD reaching ∼0.6, the mediums were transferred into 16 °C, 180 rpm incubator and protein expression was induced with 0.5 mM isopropyl β-D-1-thiogalactopyranoside (IPTG) followed by overnight growth. For GST-tagged proteins, bacteria were lysed in GST-Lysis Buffer (50 mM Tris-HCl, 600 mM NaCl, 1 mM EDTA, 1 mM DTT, 10 % Glycerol, Protease Inhibitor Cocktail, pH7.6) followed 10 minutes ultrasonic crushing and 20 minutes 12000 rpm centrifugation, and then purified using GST-Sefinose(TM) Resin (BBI) according to the manufacturer’s instructions. GST-tagged proteins were eluted by 10 mM glutathione and GST was removed by enzyme ProScission.

### *In vitro* methyltransferase assays

*In vitro* methyltransferase assays were performed using 30 ul reaction buffer (50 mM Tris-HCl pH 7.8, 50 mM KCl, 5 mM MgCl2) with 1 ul ^3^H-AdoMet (Perkin Elmer, specifific activity = 55-85 Ci/mMole,0.55 μCi/ul) or 1 mM unlabeled AdoMet for MS analysis of samples. For methylation of cell extract, reactions contained 80 µg substrates and 2 µg METTL9. For methylation of recombinant GST-tag proteins and 6×His Histone H3, reactions contained 2 µg substrates and 2 µg METTL9. Reactions were incubated at 37 °C for 1 h, and stopped by the addition of SDS-PAGE loading buffer. Proteins were separated by SDS-PAGE. For autoradiography analysis, the proteins were then transferred to PVDF membranes, stained with Ponceau S and exposed to XBT X-ray film (Carestream) for 30 days (cell extract) or 7 days (recombinant substrates). For methylation of synthetic peptides, reactions contained 5 µg substrates and 2 µg GST-METTL9 binding with GST-Sefinose (TM) Resin, GST-METTL9 was then removed by centrifugation. For autoradiography analysis, the peptides were directly loaded to NC membranes and exposed to XBT X-ray film (Carestream) for 7 days.

### Fluorescence imaging of Zn^2+^ level

Cell were cultured in the glass bottom micro-well dishes (Biosharp) for confocal microscopy imaging. Before ratiometric imaging with LSM 710 microscopy, 2 uM FluoZin-3 AM (Invitrogen) was add to dishes to stain cells for 1 hour and then was washed twice with PBS. For overexpression, the cells in 10 cm dish were infected with Slc39a7 wildtype or mutants in Plvx-IRES-BFP virus. After 24 h, the BFP-positive cells were sorted by flow cytometry. After cultured in the glass bottom micro-well dishes for 24 hours, the cells were stained using FluoZin-3 AM (2 uM) for confocal microscopy imaging.

### RNA-seq data processing and analysis

The RM-1 cells of 1×10^7^ WT and *Mettl9* KO were extracted with TRNzol (TIANGEN) for total RNA library preparation. The libraries are directly sequenced using next-generation sequencing technologies. After filtering of adaptors and low quality reads, clean reads were mapped to the mouse reference genome using HISAT/Bowtie2 tool. Mapping results were stored in BAM files using SAMtools. Total read counts at the gene level were summarized using feature-counts function in R environment, with the R package biomaRt for gene and transcript mapping. The differential expression genes were analyzed by DESeq2 with default settings using total read counts as input and the adjusted P value (p.adj) less than 0.05(39). Dotplot of differential genes of normalized expression matrix from DESeq2 analysis from GO pathway enrichment (https://wego.genomics.cn/).

We performed gene set enrichment analysis (GSEA) using KEGG function of clusterProfiler package in RStudio(40). The RNA-seq data of non-targeting control (NC) and two individual *Slc39a7* siRNA in MDA-MB-231 cells are from the accession number GSE155437 from NCBI Gene Expression Omnibus (PMID: 33608508)(34). Significant KEGG pathways with an enrichment score > 0.4 and a p-value <0.01 were identified. The R package ggplot2 was applied to visualize the results(41). GSEA revealed that the cell cycle, DNA replication, homologous recombination, ribosome biogenesis in eukaryotes, RNA transport and mismatch repair pathways showed significant enrichment in wild type samples compared with *Slc39a7* knockdown and *Mettl9* knockout samples.

### TCGA and GTEx data analysis

We download the expression data of *METTL9* in PRAD, PAAD, LIHC combing GTEx normal sample from USCS Xena (http://xena.ucsc.edu), and make boxplot with significance difference index through ggplot in R environment.

We applied ESTIMATE computational method to calculate the immune score of different cancer samples from The Cancer Genome Atlas (TCGA) database(42). Correlation between several genes expression levels and immune scores were performed using the ‘rcorr’ function implemented in Hmisc R package v.4.5.0, adopting Pearson’s correlation method. Significant correlations (P<0.05) were plotted using the pheatmap R package v.1.0.12. Genes with no notable correlation show a coefficient equal to zero.

### Statistical analysis

All experimental operations were independently repeated for at least three times. Images was analyzed using Image J Software (National Institutes of Health, Bethesda, MD, USA). Statistical analysis was performed using GraphPad Prism Software 6 (GraphPad Software, La Jolla, CA, USA). For data that had a normal distribution and homogeneity of variance, two-tailed Student’s t test was performed to evaluate significant differences between two groups. P-values<0.05 were considered statistically significant.

## Acknowledgements

We thank Yong Ding for providing with the plasmids GST-SDG714C and PET-28a-Histone H3. We thank the proteomics platform and imaging platform of core facility center for life sciences, USTC. We thank the platform of autoradiography in core facility center for life sciences and Xuebiao Yao lab, USTC. We thank Shu Zhu for useful discussion. This work was supported by NSFC grant 81872327 (to WP), “USTC Important Direction” Cultivation Project (WK3520000013) (to WP), “USTC New Medicine” Cultivation Project (WK9110000065) (to DC)

## Author Contributions

ML, DC and WP. designed the experiments. ML, DC performed and interpreted the experiments. CH, SL collected all MS data. LZ, PZ, and LZ performed the bio-information analyses; AY, ZY, CLand KZ provided critical comments and suggestions; WP, DC and ML wrote the manuscript; WP and DC supervised the project. The authors declare no competing interests.

## References

1. Clarke S. Protein methylation. Current opinion in cell biology. 1993;5(6):977–83.

2. Kwiatkowski S, Drozak J. Protein histidine methylation. Current protein & peptide science. 2020.

3. Bannister AJ, Schneider R, Kouzarides T. Histone methylation: Dynamic or static? Cell. 2002;109(7):801–6.

4. Michalak EM, Burr ML, Bannister AJ, Dawson MA. The roles of DNA, RNA and histone methylation in ageing and cancer. Nature reviews Molecular cell biology. 2019;20(10):573–89.

5. Jambhekar A, Dhall A, Shi Y. Roles and regulation of histone methylation in animal development. Nature reviews Molecular cell biology. 2019;20(10):625–41.

6. Biggar KK, Wang Z, Li SS. SnapShot: Lysine Methylation beyond Histones. Mol Cell. 2017;68(5):1016–e1.

7. Di Blasi R, Blyuss O, Timms JF, Conole D, Ceroni F, Whitwell HJ. Non-Histone Protein Methylation: Biological Significance and Bioengineering Potential. ACS Chem Biol. 2021;16(2):238–50.

8. Rodriguez-Paredes M, Lyko F. The importance of non-histone protein methylation in cancer therapy. Nature reviews Molecular cell biology. 2019;20(10):569–70.

9. Searle JM, Westall RG. The Occurrence of Free Methylhistidine in Urine. Biochemical Journal. 1951;48(5):R50–R.

10. Kwiatkowski S, Seliga AK, Vertommen D, Terreri M, Ishikawa T, Grabowska I, et al. SETD3 protein is the actin-specific histidine N-methyltransferase. eLife. 2018;7.

11. Guo Q, Liao S, Kwiatkowski S, Tomaka W, Yu H, Wu G, et al. Structural insights into SETD3-mediated histidine methylation on beta-actin. eLife. 2019;8.

12. Wilkinson AW, Diep J, Dai S, Liu S, Ooi YS, Song D, et al. SETD3 is an actin histidine methyltransferase that prevents primary dystocia. Nature. 2019;565(7739):372–6.

13. Al-Hadid Q, Roy K, Munroe W, Dzialo MC, Chanfreau GF, Clarke SG. Histidine methylation of yeast ribosomal protein Rpl3p is required for proper 60S subunit assembly. Molecular and cellular biology. 2014;34(15):2903–16.

14. Webb KJ, Zurita-Lopez CI, Al-Hadid Q, Laganowsky A, Young BD, Lipson RS, et al. A Novel 3-Methylhistidine Modification of Yeast Ribosomal Protein Rpl3 Is Dependent upon the YIL110W Methyltransferase. Journal of Biological Chemistry. 2010;285(48):37598–606.

15. Ning Z, Star AT, Mierzwa A, Lanouette S, Mayne J, Couture JF, et al. A charge-suppressing strategy for probing protein methylation. Chemical communications. 2016;52(31):5474–7.

16. Dai S, Horton JR, Wilkinson AW, Gozani O, Zhang X, Cheng X. An engineered variant of SETD3 methyltransferase alters target specificity from histidine to lysine methylation. The Journal of biological chemistry. 2020;295(9):2582–9.

17. Bokar JA, Shambaugh ME, Polayes D, Matera AG, Rottman FM. Purification and cDNA cloning of the AdoMet-binding subunit of the human mRNA (N6-adenosine)-methyltransferase. Rna. 1997;3(11):1233–47.

18. Liu J, Yue Y, Han D, Wang X, Fu Y, Zhang L, et al. A METTL3-METTL14 complex mediates mammalian nuclear RNA N6-adenosine methylation. Nature chemical biology. 2014;10(2):93–5.

19. Pendleton KE, Chen B, Liu K, Hunter OV, Xie Y, Tu BP, et al. The U6 snRNA m(6)A Methyltransferase METTL16 Regulates SAM Synthetase Intron Retention. Cell. 2017;169(5):824–35 e14.

20. Jiang X, Liu B, Nie Z, Duan L, Xiong Q, Jin Z, et al. The role of m6A modification in the biological functions and diseases. Signal Transduct Target Ther. 2021;6(1):74.

21. Malecki J, Jakobsson ME, Ho AYY, Moen A, Rustan AC, Falnes PO. Uncovering human METTL12 as a mitochondrial methyltransferase that modulates citrate synthase activity through metabolite-sensitive lysine methylation. The Journal of biological chemistry. 2017;292(43):17950–62.

22. Shimazu T, Barjau J, Sohtome Y, Sodeoka M, Shinkai Y. Selenium-based S-adenosylmethionine analog reveals the mammalian seven-beta-strand methyltransferase METTL10 to be an EF1A1 lysine methyltransferase. PloS one. 2014;9(8):e105394.

23. Rhein VF, Carroll J, Ding SJ, Fearnley IM, Walker JE. Human METTL12 is a mitochondrial methyltransferase that modifies citrate synthase. FEBS letters. 2017;591(12):1641–52.

24. Woodcock CB, Yu D, Hajian T, Li J, Huang Y, Dai N, et al. Human MettL3-MettL14 complex is a sequence-specific DNA adenine methyltransferase active on single-strand and unpaired DNA in vitro. Cell discovery. 2019;5(1).

25. Roy A, Kucukural A, Zhang Y. I-TASSER: a unified platform for automated protein structure and function prediction. Nat Protoc. 2010;5(4):725–38.

26. Yang J, Yan R, Roy A, Xu D, Poisson J, Zhang Y. The I-TASSER Suite: protein structure and function prediction. Nat Methods. 2015;12(1):7–8.

27. Yang J, Zhang Y. I-TASSER server: new development for protein structure and function predictions. Nucleic acids research. 2015;43(W1):W174–81.

28. Raftery MJ, Harrison CA, Alewood P, Jones A, Geczy CL. Isolation of the murine S100 protein MRP14 (14 kDa migration-inhibitory-factor-related protein) from activated spleen cells: Characterization of post-translational modifications and zinc binding. Biochemical Journal. 1996;316:285–93.

29. Carroll J, Fearnley IM, Skehel JM, Runswick MJ, Shannon RJ, Hirst J, et al. The post-translational modifications of the nuclear encoded subunits of complex I from bovine heart mitochondria. Molecular & Cellular Proteomics. 2005;4(5):693–9.

30. Bin BH, Seo J, Kim ST. Function, Structure, and Transport Aspects of ZIP and ZnT Zinc Transporters in Immune Cells. J Immunol Res. 2018;2018:9365747.

31. Ressnerova A, Raudenska M, Holubova M, Svobodova M, Polanska H, Babula P, et al. Zinc and Copper Homeostasis in Head and Neck Cancer: Review and Meta-Analysis. Current medicinal chemistry. 2016;23(13):1304–30.

32. Sanna A, Firinu D, Zavattari P, Valera P. Zinc Status and Autoimmunity: A Systematic Review and Meta-Analysis. Nutrients. 2018;10(1).

33. Woodruff G, Bouwkamp CG, de Vrij FM, Lovenberg T, Bonaventure P, Kushner SA, et al. The Zinc Transporter SLC39A7 (ZIP7) Is Essential for Regulation of Cytosolic Zinc Levels. Molecular pharmacology. 2018;94(3):1092–100.

34. Chen PH, Wu JL, Xu YT, Ding CKC, Mestre AA, Lin CC, et al. Zinc transporter ZIP7 is a novel determinant of ferroptosis. Cell death & disease. 2021;12(2).

35. Zhang T, Liu J, Fellner M, Zhang C, Sui D, Hu J. Crystal structures of a ZIP zinc transporter reveal a binuclear metal center in the transport pathway. Science advances. 2017;3(8):e1700344.

36. Zhang T, Kuliyev E, Sui D, Hu J. The histidine-rich loop in the extracellular domain of ZIP4 binds zinc and plays a role in zinc transport. The Biochemical journal. 2019;476(12):1791–803.

37. Adulcikas J, Norouzi S, Bretag L, Sohal SS, Myers S. The zinc transporter SLC39A7 (ZIP7) harbours a highly-conserved histidine-rich N-terminal region that potentially contributes to zinc homeostasis in the endoplasmic reticulum. Computers in biology and medicine. 2018;100:196–202.

38. Davydova E, Shimazu T, Schuhmacher MK, Jakobsson ME, Willemen H, Liu T, et al. The methyltransferase METTL9 mediates pervasive 1-methylhistidine modification in mammalian proteomes. Nature communications. 2021;12(1):891.

39. Love MI, Huber W, Anders S. Moderated estimation of fold change and dispersion for RNA-seq data with DESeq2. Genome biology. 2014;15(12).

40. Yu G, Wang LG, Han Y, He QY. clusterProfiler: an R package for comparing biological themes among gene clusters. OMICS. 2012;16(5):284–7.

41. Villanueva RAM, Chen ZJ. ggplot2: Elegant Graphics for Data Analysis, 2nd edition. Meas-Interdiscip Res. 2019;17(3):160–7.

42. Yoshihara K, Shahmoradgoli M, Martinez E, Vegesna R, Kim H, Torres-Garcia W, et al. Inferring tumour purity and stromal and immune cell admixture from expression data. Nature communications. 2013;4:2612.

