## Supplementary figures for "METTL9 regulates N1-histidine methylation of zinc transporters to promote tumor growth"

Figure S1

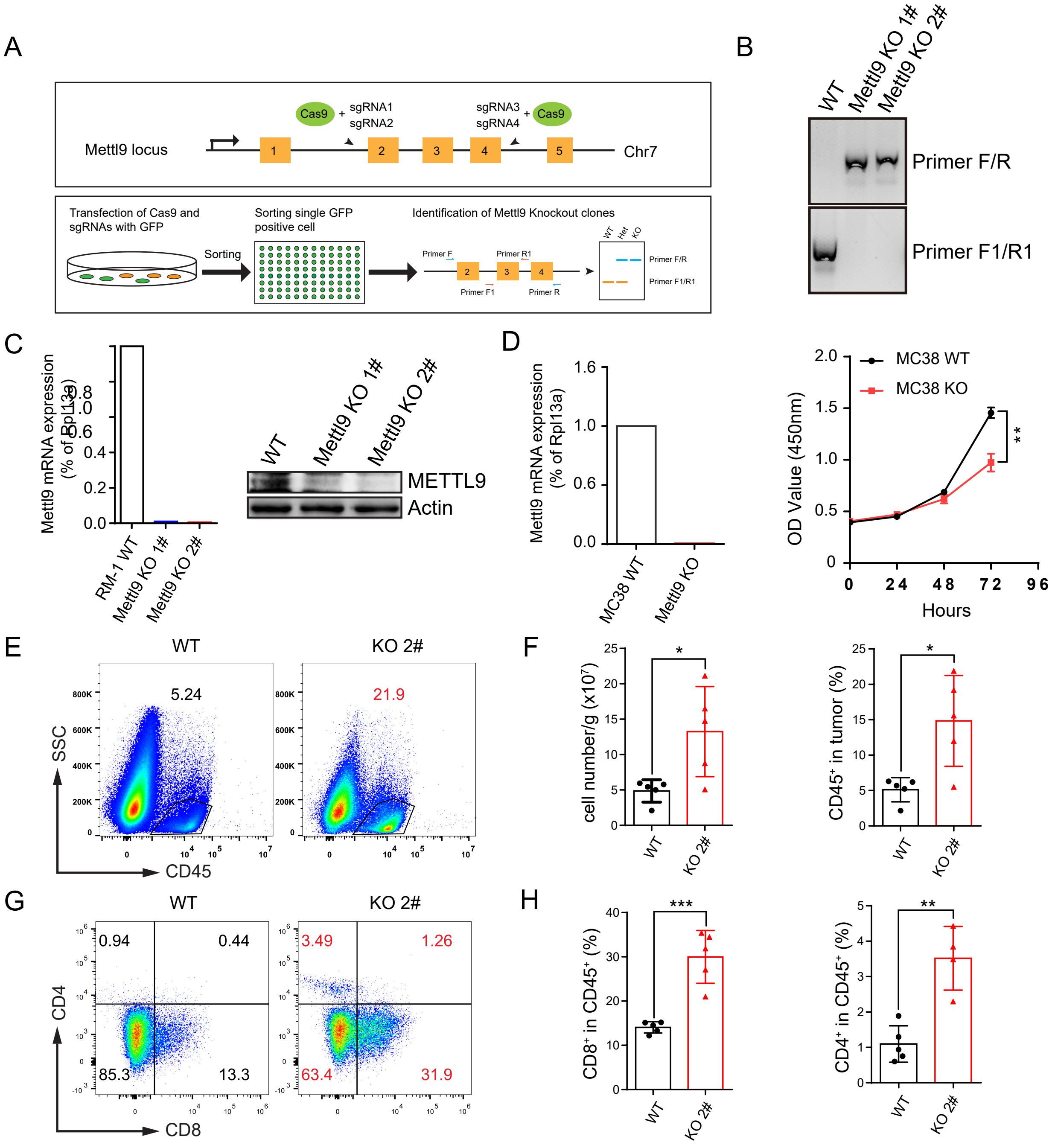


**Figure S1. *Mettl9* is required for cell proliferation and tumor growth.**

**(A)** The scheme of constructing *Mettl9* knockout cell line. CRISPR/Cas9 is used to remove exon 2 and 3 and exon 4 of *Mettl9*. **(B, C)** Validation of the efficiency of *Mettl9* knockout by measurement of mRNA and protein expression. **(D)** Knockout of *Mettl9* in MC38 tumor cells decreases cell growth measured by CCK-8 assay. **(E-H)** WT and *Mettl9* KO RM-1 tumor cells were injected into C57B6/J mouse (n=5). Tumor infiltrating immune cells from WT and *Mettl9* KO tumor-bearing C57B6/J mice were examined by flow cytometry. **(E, F)** The cell numbers per tumor weight and the percentages of tumor-infiltrating $CD45^+$ cells within all live cells from the tumor were quantified. Each dot represents one mouse. n=5. **(G, H)** Tumor-infiltrating $CD4^+$ and $CD8^+$ T cells were quantified as the percentages within total $CD45^+$ cells. Each dot represents one mouse. n=5. For all panels, *: $P< 0.05$; **: $P < 0.01$; ***: $P < 0.001$; ****: $P < 0.0001$. Error bars represent S.D. Data are representative of three independent experiments.

Figure S2

A

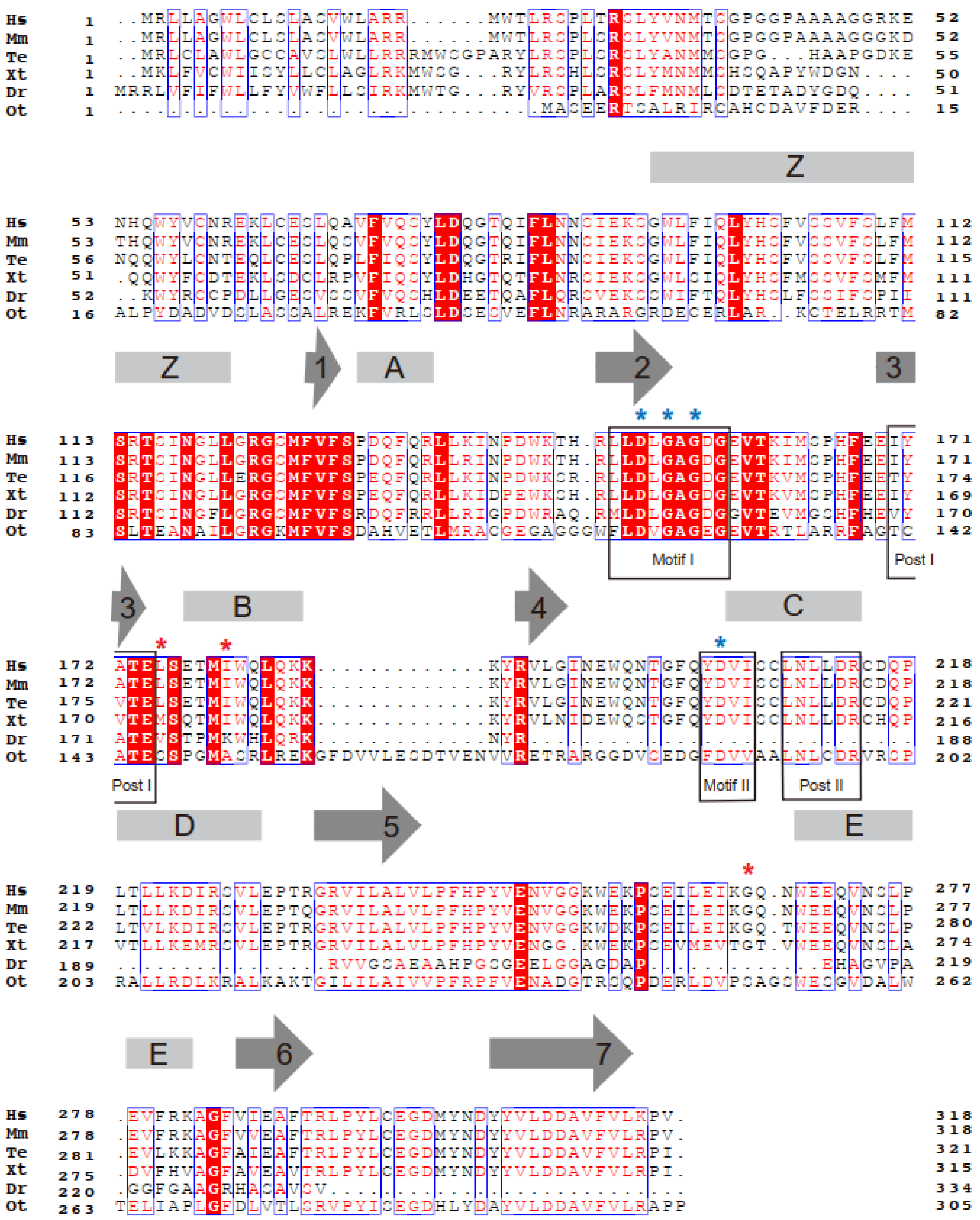

**Figure S2. METTL9 is a conservative 7-beta Strand methyltransferase.**

**(A)** Alignment of the core methyltransferase domain of METTL9 orthologues. Homo Sapiens (Hs; NP_057109.3), Mus musculus (Mm; NP_067529.2), Thamnophis elegans (Te; XP_032087096.1), Xenopus tropicalis (Xt; NP_001007899.1), Danio rerio (Dr; NP_001070810.1), Ostreococcus tauri (Ot; XP_003078360.1). Blue star, mutant at predicted motifs. Red star, mutant at predicted active sites.

Figure S3

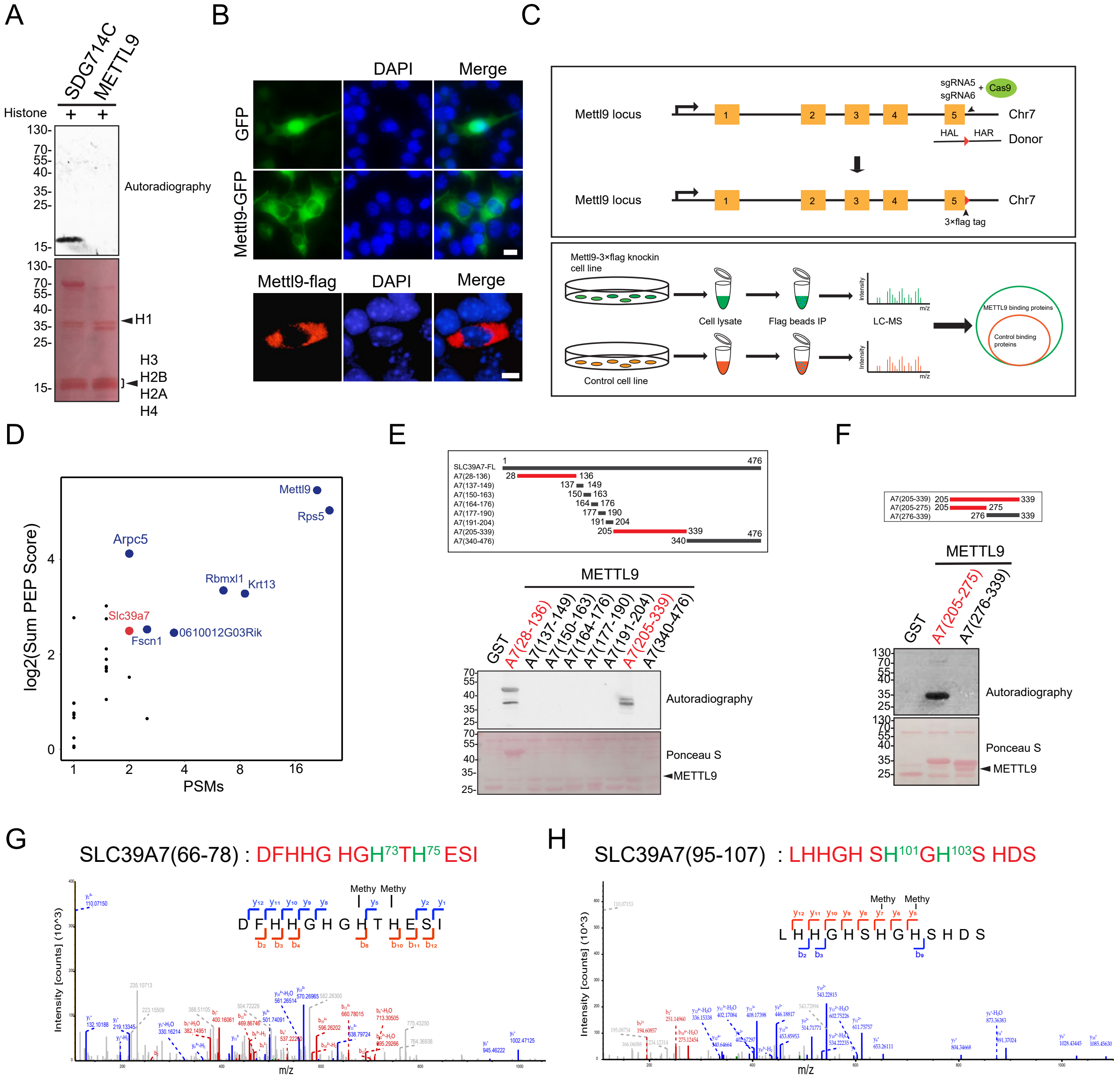


**Figure S3. METTL9 methylates $Zn^{2+}$ transporter SLC39A7.**

**(A)** Fluorography showing histone was not methylated by METTL9. GST-SDG714C as a positive control. **(B)** Immunofluorescence localization analysis of *Mettl9*-GFP fusion protein in B16F10 cell (exogenous) and *Mettl9*-flag fusion protein in RM-1 *Mettl9*-3×flag knock-in cells (endogenous). Scale bar, 10 μm. **(C)** The scheme of constructing flag-tag knock-in RM-1 tumor cell line and the experimental workflow. **(D)** The METTL9 binding proteins. Dotplot of METTL9-interacting proteins by immunoprecipitation–mass spectrometry (IP–MS) analysis. Colored indicates enriched proteins compared to control (PSMs > 2; log2(Sum PEP Score) > 2). **(E, F)** Fine mapping of the METTL9-methylated regions in SLC39A7. The diagram shows different recombinant GST-SLC39A7 (Termed A7) truncates. The methylated peptide was colored in red. **(G, H)** LC-MS/MS fragmentation spectra analysis of METTL9 methylated peptide SLC39A7(66-78) and SLC39A7(95-107). The monomethylation residues at His73, His75, His101 and His103 were colored in green.

FigureS4

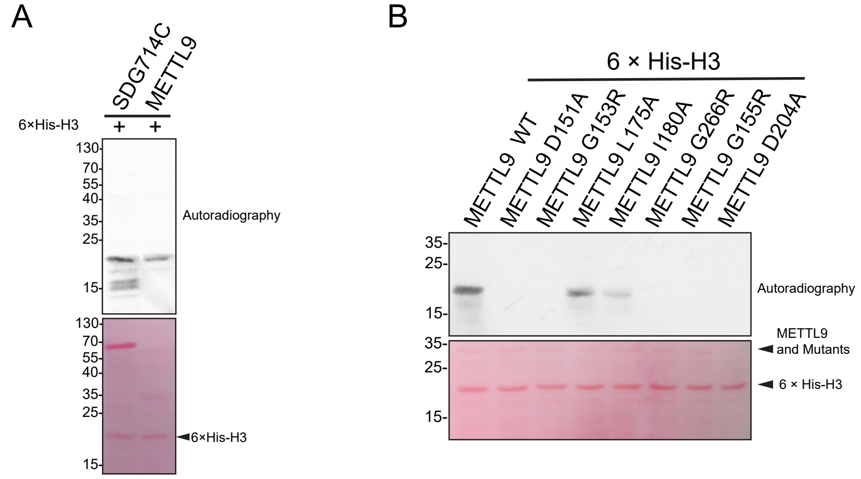


**Figure S4. The key sites of METTL9 enzymatic activity**

**(A)** Fluorography showing recombinant histone 3 with 6×His tag was methylated by METTL9. GST-SDG714C as a positive control. Ponceau S-stained membrane as loading control (bottom)
(**B**) *In vitro* activity of WT and mutated METTL9 on recombinant 6×His-histone H3. D151A, G153R, G155R, in motif-I; D204A, in motif-II; L175A, I180A, G266R, predicted active sites. The data are representative of three independent experiments

Figure S5

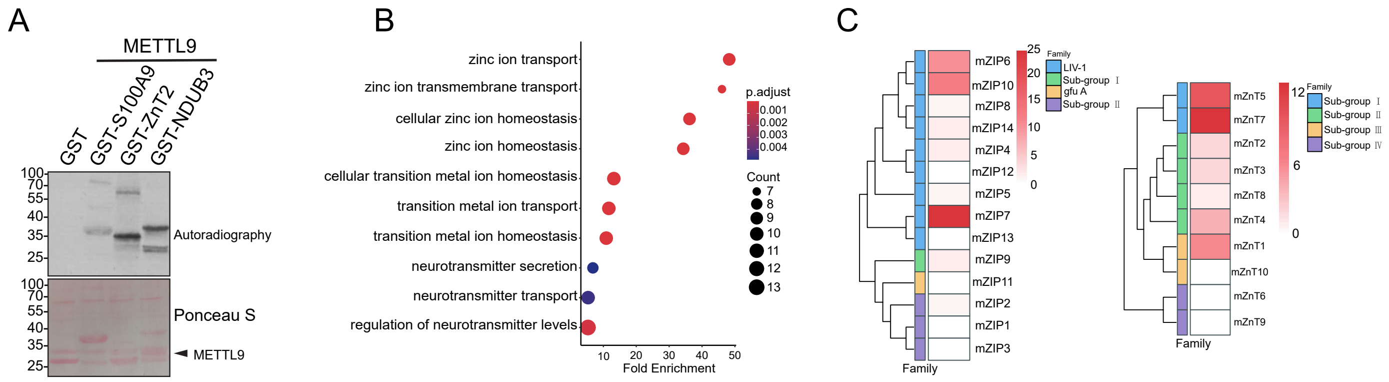


**Figure S5. METTL9 methylates xHxH motif on zinc transporters.**

**(A)** *In vitro* activity of METTL9 on recombinant GST-S100A9, GST-ZnT2 and GST-NDUB3 with GHxH motif. GST as a control substrate. **(B)** GO analysis of top pathways from proteins enriched with GHA/C/G/SH motif. **(C)** Evolutionary tree enrichment of proteins containing GHA/C/G/SH motif of ZnT (SLC30s) and ZIP (SLC39s) family.

Figure S6

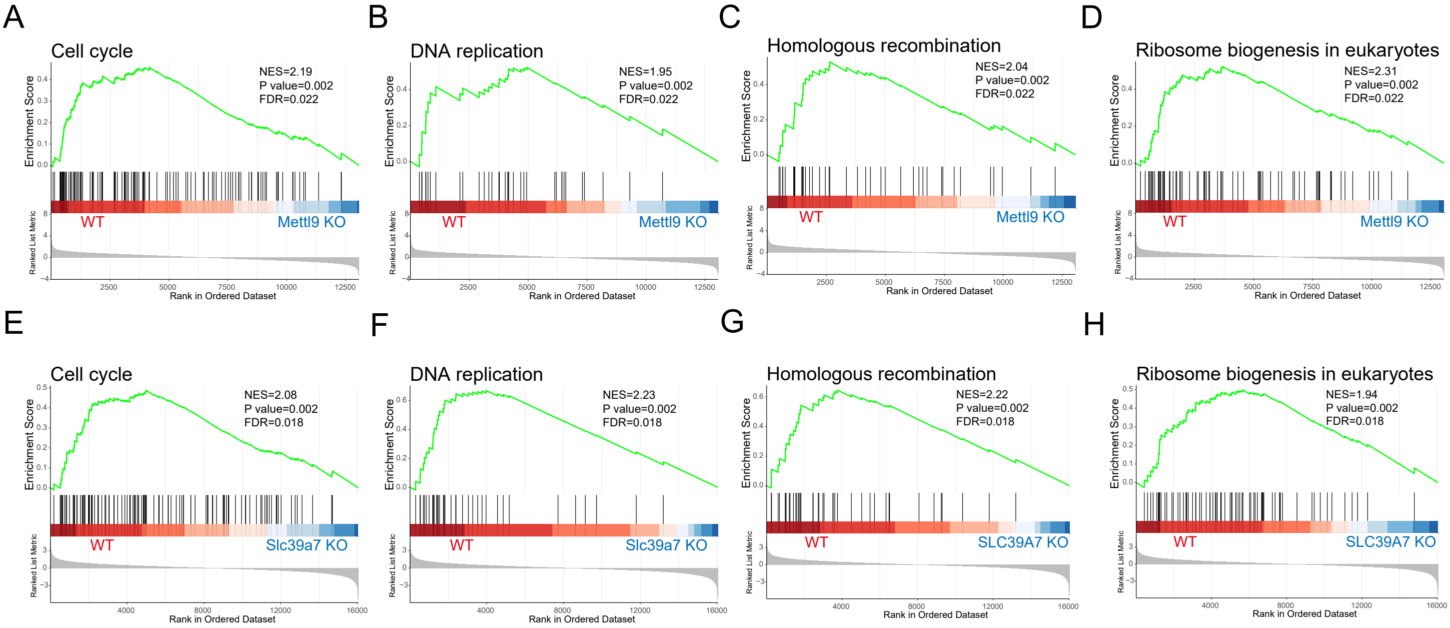


**Figure S6. *Mettl9* and *Slc39a7* regulates common downstream pathways**

**(A-H)** Gene set enrichment analysis (GSEA) of differentially expressed genes in *Mettl9* KO cell or *Slc39a7* KD cells. Significant KEGG pathways with an enrichment score > 0.4 and p-value <0.01 were identified. Transcripts downregulated in *Mettl9* KO cell or *Slc39a7* KD cell versus their controls significantly enriched in **(A, E)** Cell cycle, **(B, F)** DNA replication, **(C, G)** Homologous recombination and **(D, H)** Ribosome biogenesis in eukaryotes.

Figure S7

A

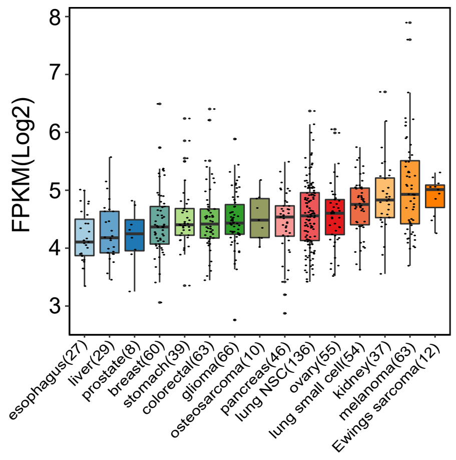


**Figure S7.** ***METTL9*** **transcript levels were analyzed in cancer cell lines in CCLE.**
